# Caloric restriction and intermittent fasting during lactation are linked to impaired maternal care, increased impulsivity and amygdala redox imbalance in dams

**DOI:** 10.64898/2026.07.03.736282

**Authors:** Nícollas Costa Veloso, Raul Carvalho Dayrell, Larissa Neves Roque, Sofia Veloso Duarte, Mateus Teixeira Lages Santos, Victor Emanuel dos Reis Advíncola, Alexandre Alves da Silva, Nísia Andrade Villela Dessimoni Pinto, Valentina Mosienko, Arthur Rocha-Gomes, Tania Regina Riul

## Abstract

The lactational period requires substantial metabolic and behavioral adaptations, and more than 70% of mothers report weight concerns and attempt weight loss by four months postpartum. Nevertheless, how distinct restrictive paradigms during lactation alter maternal behavior, and the extent to which associated neurochemical changes modulate these behaviors, remains poorly understood.

In the current study, we modeled restrictive diets in lactating rats to evaluate caregiving behavior and its relationship to amygdalar redox status. Intermittent fasting (IF) and caloric restriction (CR) administered to lactating Wistar dams from postpartum day 0 to day 28 impaired maternal care, evidenced by delayed pup retrieval, reduced nest building, and decreased nursing frequency relative to *ad libitum*-fed controls. Both diets reduced body and adipose tissue weight, and energy efficiency. IF and CR increased impulsivity-like phenotype: CR doubled open-arm exploration in the elevated plus maze; IF and CR increased center-zone exploration in the open field by three- and two-fold, respectively; IF doubled time in the light-dark box light compartment. A composite maternal behavioral score showed impairment in dams in both IF and CR groups. At the neurochemical level, both diets reduced amygdalar superoxide dismutase activity, which correlated negatively with the maternal behavioral score.

Both restrictive diets produced an underweight phenotype with weakened dam-pup interactions and increased impulsivity. These behavioral changes co-occurred with amygdalar redox imbalance, which correlated with the severity of maternal impairment. Overall, the study refines understanding of the nutritional and behavioral consequences of dietary restriction in lactation and implicates disrupted redox homeostasis as a plausible mechanism.

## 1 INTRODUCTION

The lactational period, characterised by the arrival of the offspring, is a phase of intense transformation in a woman’s life. Among the common concerns during this time, weight gain acquired during pregnancy emerges as an important issue, often driven by social pressures on mothers to return rapidly to their pre-pregnancy body shape. For example, 75% of mothers report concerns about their weight and body image one month after delivery, and by four months postpartum, 70% engage in some form of weight-loss attempt (Martínez-Olcina et al., 2020). Consequently, some women consider adopting restrictive diets as a strategy to achieve this goal, and it has been reported that the occurrence of eating disorders during this period may triple compared to pregnancy (Pettersson; Zandian; Clinton, 2016).

It is important to understand that weight gain is a natural physiological response to the demands that will arise during breastfeeding and offspring care. The body undergoes hormonal changes, accumulating essential reserves to sustain both the mother and offspring during a phase of increased energy expenditure, including the onset of milk production, lactation, and maternal care (Muro-Valdez et al., 2023). In line with these considerations, a balanced maternal diet during lactation also influences both the quantity and quality of breast milk, whereas dietary restriction may compromise maternal care, breastfeeding, and neonatal health (Jawaid; Jehle; Mansuy, 2021; Öner Sayar; Köseoğlu, 2025).

Evidence indicates that maternal nutrient deficiency or lack of access to a balanced diet is directly associated with reduced caregiving quality, for example, in the frequency and adequacy of breastfeeding, thereby impairing offspring development (Frith et al., 2012; Prado et al., 2018; Tomaszewska et al., 2025). Maternal mental health may also be affected, as a restrictive diet, combined with the complexity of motherhood, may increase the risk of postpartum anxiety and/or depression (Walker-Mao et al., 2024). Among the restrictive diets adopted by lactating women, intermittent fasting (IF) and caloric restriction (CR) have emerged as popular trends and may negatively affect both maternal health and offspring development.

Intermittent fasting (IF) consists of alternating periods of feeding and complete abstinence from food. Among the protocols adopted are the 16:8 model (16 hours of fasting and 8 hours of feeding) and alternate-day fasting, with 24 hours of food deprivation interspersed with 24 hours of *ad libitum* feeding (Enríquez Guerrero et al., 2021). The latter is considered more drastic and has been successfully used in rodent models to investigate metabolic adaptations, redox status, and behaviour in the pups (Rocha-Gomes et al., 2026a). Another model of food restriction is CR, characterised by a 50% reduction in daily dietary intake, which has been widely employed in animal models to examine its effects on offspring growth, physiology, metabolism, and behaviour (Rocha-Gomes et al., 2021a). However, most available studies have focused primarily on the effects of these diets on offspring physiology, whereas investigations involving IF diet and those centred on maternal outcomes remain scarce.

One metabolic aspect that may be compromised by restrictive diets is the redox status of multiple tissues, which refers to the dynamic balance between reactive species production and the organism’s antioxidant capacity. Notably, studies in both humans (Catal et al., 2007; Jain; Jadhav; Varma, 2013) and rodents (Gonçalves et al., 2026a) demonstrate that restrictive diets can substantially reduce antioxidant defenses, thereby impairing the function of several organs. For example, Sinha et al. (2020) showed that protein restriction during the first 15 days of lactation in Sprague-Dawley rats induced redox imbalance in the hippocampus of male and female offspring during adolescence and adulthood. In addition, our group showed that both caloric restriction and intermittent fasting during lactation alter superoxide dismutase (SOD) activity in the amygdala of adolescent male and female offspring (Rocha-Gomes et al., 2026a), a brain region critically involved in stress and anxiety responses (Tovote; Fadok; Lüthi, 2015). Consequently, during periods of high energetic demand, such as lactation, this type of dietary restriction may also exert harmful effects on the mother, representing a potential mechanistic pathway linking lactational malnutrition to metabolic and mental health outcomes.

However, it remains unclear whether restrictive dietary conditions during lactation can alter maternal care and behaviour in association with redox alterations in the amygdala. Therefore, the present study exposed Wistar rat dams to two restrictive dietary restrictions, IF and CR, during lactation and aimed to evaluate nutritional status, maternal caregiving and dam-pup interactions, anxiety-like behaviour, and amygdalar redox balance.

## 2 MATERIAL AND METHODS

### 2.1 Animals and Ethics

This study was conducted at the Experimental Nutrition Laboratory (LabNutrex) of the Federal University of the Jequitinhonha and Mucuri Valleys (UFVJM). The experiment was carried out in accordance with the ethical principles of the *National Institutes of Health Guide for the Care and Use of Laboratory Animals* (NIH Publications No. 80-23) and was approved by the Institutional Animal Care and Use Committee (CEUA-UFVJM – protocol 055/2019).

Animals were obtained from LabNutrex, underwent a two-week acclimatisation period before the experiment, and were housed in polypropylene cages (41 × 34 × 16 cm) placed on ventilated racks (Alesco^®^), under standard conditions (natural humidity; controlled air renewal; temperature 22°C ± 2; 12-hour light/dark cycle, with lights on at 06:00 a.m.) and had free access to potable water.

### 2.2 Experimental design

Male and nulliparous female Wistar rats (*Rattus norvegicus*), aged 70 days at the start of the experiment, were used for polygamous mating (two females per male) for two weeks. After the gestation period, litters were standardised at birth (postnatal day 0; PND0) to consist of one dam and eight pups (4 males and 4 females). To minimise genetic factors, pups born on the same day were cross-fostered among litters, a procedure routinely used in our laboratory in which dams do not reject the fostered pups (Rocha-Gomes et al., 2021b, 2024). Subsequently, the dams were randomly assigned to receive one of the following diets, from PND0 until PND28: Control (C) – dams received standard chow (Nuvilab CR-1^®^) *ad libitum* (n = 12 litters); Intermittent Fasting (IF) – dams received standard chow *ad libitum* for the first 24 hours, followed by a 24-hour period of without access to food, alternating feeding and restriction periods every 24 hours (n = 11 litters); Caloric Restriction (CR) – dams received 50% of standard chow intake consumed by control dams (n = 11 litters).

The IF and CR methodologies followed the protocols previously executed and described by our group (Gonçalves et al., 2026b; Rocha-Gomes et al., 2026b). The present study evaluated the dams; all pups were assigned to another experiment on weaning (PND21). Figure 1 illustrates the study’s experimental design.

**Figure 1.**
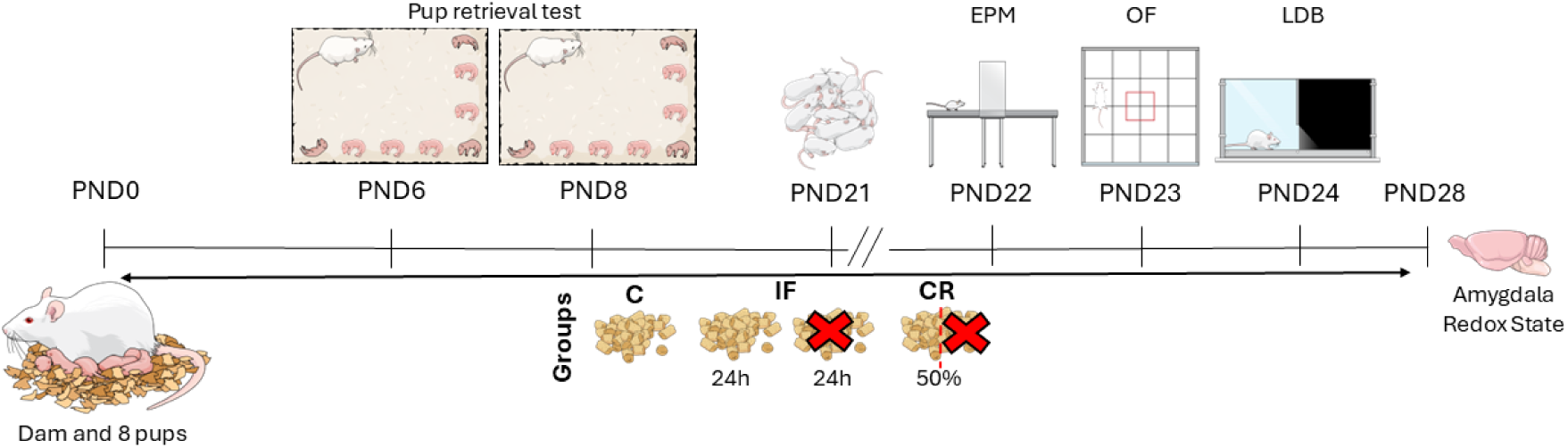
Schematic illustration of the experimental design. Litters were formed on postnatal day 0 (PND0), each consisting of one dam and eight pups (4 males and 4 females). Between PND0 and PND28, dams received one of the following diets: Control (C), intermittent fasting (IF), or caloric restriction (CR). The pup retrieval test was conducted on PND6 and PND8. Offspring were weaned on PND21 and were not included in this study. Between PND22 and PND24, dams underwent the elevated plus maze (EPM), open field (OF), and light-dark box (LDB) tests, respectively. On PND28, dams were euthanised, and brain tissue (amygdala) was collected for redox analysis. Figure created in the Mind the Graph platform.

### 2.3 Nutritional Assessments

Maternal food intake was recorded daily between 12:00 and 14:00 using a semi-analytical balance, and total intake was calculated as the cumulative consumption from PND0 to PND28. Caloric intake was estimated by multiplying the total amounts of carbohydrates, lipids, and proteins consumed by the factors 4, 9, and 4, respectively. Maternal body weight was measured individually once weekly during sawdust replacement. From these data, weight variation (PND28-PND0) and energy efficiency (weight gain/energy intake) were calculated (Da Silva et al., 2023).

On PND28, dams were anaesthetised with ketamine (60 mg.kg⁻¹) and xylazine (15 mg.kg⁻¹), and body length as well as abdominal (AC) and thoracic (TC) circumferences were measured to calculate body mass index (BMI = weight/body length²) and the AC/TC ratio. Subsequently, dams were euthanised by decapitation, and blood, brain tissue, organs (spleen, heart, liver, kidneys, and adrenal glands), and abdominal adipose tissue (visceral, retroperitoneal, mesenteric, and gonadal) were collected for analysis. Organs and adipose tissues were immersed in saline solution, cleaned, blotted dry on filter paper, weighed using an analytical balance (Shimadzu^®^), and stored at -80°C until further processing (Da Silva et al., 2023)

Approximately 2 mL of blood was collected, and serum was obtained for the determination of total cholesterol, high-density lipoprotein (HDL), triacylglycerol, aspartate aminotransferase (AST), creatinine, and urea by spectrophotometry, following the manufacturer’s instructions (Labtest^®^) (Da Silva et al., 2023).

### 2.4 Behavioural Assessments

All behavioural tests were conducted in the LabNutrex behavioural testing room by an experienced researcher. The room was equipped with an exhaust fan and a fluorescent lamp (100 lux illumination). All procedures were performed in the morning (07:00-12:00), under double-blind conditions, with the researcher absent during video recording (Sony^®^ Handycam). The apparatus used for behavioural testing was cleaned with 70% ethanol between animals to eliminate olfactory cues.

#### 2.4.1 Pup Retrieval Test

Pup retrieval test was performed on PND6 and PND8 to evaluate maternal motivation, attentional engagement, and efficiency in pup retrieval, regrouping, and caregiving following brief separation and nest disruption (Riul et al., 1999a).

For testing on both days, the dams were first removed from the box, and the pups were distributed along their sides. The dam was then returned to her home cage, and behaviour was recorded for 10 minutes. The following maternal parameters were analysed: latency to retrieve the first pup (time between returning the dam to the box and picking up the first pup with the mouth); latency to nest building (time between returning the dam and grouping all pups in the nest); maternal preference for retrieving a male or female pup first; maternal care (dam observed touching, sniffing, licking pups, or simply remaining over them); and nursing frequency and duration (dam observed nursing the litter or at least one pup observed suckling). Non-maternal behaviours were also recorded: locomotion time (movement within the box without contact with pups) and rearing number (dam observed in an upright position supported on the hind limbs). If the dam did not retrieve any pup during the total duration of the test (10 minutes), it was excluded from this parameter (Riul et al., 1999a).

#### 2.4.2 Elevated Plus Maze (EPM)

The EPM test was conducted in dams after weaning (PND22). The EPM is based on rodents’ innate aversion to open and elevated spaces; when exposed to the apparatus, they exhibit a conflict between the drive to explore a novel environment and their aversion to the open arms (Rocha-Gomes et al., 2021b). The EPM consisted of a wooden apparatus elevated 50 cm above the floor, comprising two closed arms (50 × 10 × 40 cm; 40 lux) perpendicular to two open arms (50 × 10 cm; 100 lux) and a central area (10 × 10 cm). The open arms were fitted with 1-cm acrylic edges to prevent falls. Each dam was placed in the central area facing one of the closed arms, and behaviour was recorded for 5 minutes. The percentage of entries and time spent in the open arms, frequency of visits to the distal ends of the open arms, and rearing behaviour were analysed (Rocha-Gomes et al., 2021b).

#### 2.4.3 Open Field (OF)

The OF test was conducted on PND23 to assess locomotor/exploratory activity, and it can also reflect anxiety-like behaviour through the measurement of the frequency of entries and time spent in the central zone (Rocha-Gomes et al., 2021b). The apparatus consisted of a wooden box (70 × 70 × 50 cm), with light grey walls and floor, marked with 16 equal squares (17.5 × 17.5 cm each), with the four central squares defined as the centre zone (100 lux). Each dam was placed individually in the centre of the OF, and behaviour was recorded for 5 minutes. The variables analysed were latency to leave, frequency of entries and time spent in the centre, and number of squares crossed (distance covered).

#### 2.4.4 Light-Dark Box (LDB)

The LDB was conducted on PND24. This test is based on rodents’ innate aversion to illuminated spaces. Rodents preferentially remain longer in the dark chamber, whereas those that spend more time in the light chamber are classified as exhibiting lower anxiety-like behaviour (Rocha-Gomes et al., 2021b). The apparatus consisted of two square chambers (30 × 30 cm). The first (light chamber) had white walls and floor and was illuminated by a white lamp (100 lux). The second (dark chamber) had black walls and floor and was illuminated by a red lamp (20 lux). The chambers were connected by a rectangular opening (10 cm height), allowing free passage between them. Each dam was placed in the light chamber facing the rectangular opening, and behaviour was recorded for 5 minutes. The following parameters were assessed: frequency of entries and time spent in the light chambers; number of squares crossed in light chamber (distance covered); and frequency of rearing (Rocha-Gomes et al., 2021b).

#### 2.4.5 Maternal behavioural score

To generate an integrated index of maternal behaviour, we applied the z-score approach, a simple mathematical tool widely used to standardise behavioural outcomes across different tests, enabling the generation of an overall “emotionality index” or, in the present study, a maternal behavioural score. Z-scores were calculated by subtracting the mean of the control group from each individual value of the experimental groups and dividing the result by the corresponding standard deviation of the control group. Thus, z-scores indicate how many standard deviations an observation lies above or below the control mean (Guilloux et al., 2011). For this analysis, the following variables were selected: latency to retrieve the first pup on PND6 and PND8; time spent in the open arms of the EPM, time spent in the centre zone of the OF; and time spent in the light chamber of the LDB. These parameters were selected because they represent widely used behavioural dimensions in neuroscience (Riul et al., 1999a; Rocha-Gomes et al., 2026b), with higher scores indicating reduced maternal motivation and attentional engagement.

### 2.5 Redox Status

Brains were removed, and the amygdala from both hemispheres was dissected, homogenised in ice-cold phosphate-buffered saline (50 mM; pH 7.0; 4°C), centrifuged (4°C; 750 g; 10 minutes) and the supernatant was used for the following analyses.

Total antioxidant capacity was assessed using the ferric reducing antioxidant power (FRAP) assay, based on the reduction of ferric-tripyridyltriazine to ferrous tripyridyltriazine, measured at 550 nm, and expressed as nM FeSO₄/mg protein. Superoxide dismutase (SOD) activity was evaluated by measuring the inhibition of pyrogallol auto-oxidation at 420 nm and expressed as U/mg protein. Catalase (CAT) activity was determined by hydrogen peroxide metabolism and expressed as ΔE/min/mg protein. Lipid peroxidation was measured using the thiobarbituric acid reactive substances (TBARS) assay, with results expressed as nmol malondialdehyde (MDA)/mg protein. All redox analyses were performed in triplicate using a UV/Visible U-200 L spectrophotometer, and values were normalised to protein concentration determined by the Bradford method using bovine serum albumin (BSA) as standard (Bradford, 1976; Rocha-Gomes et al., 2021b).

### 2.6 Statistical Analysis

The normality of the data distribution was assessed using the Shapiro Wilk test, and variables with non-normal distributions were subjected to logarithmic transformation. Maternal body weight was analysed using a repeated-measures analysis of variance (ANOVA), with diet (three levels: C, IF, and CR) as the between-subjects factor, and time (four levels: 0, 7, 14, 21, and 28 days) as the repeated-measures factor. All other maternal data were analysed using one-way ANOVA with diet (three levels: C, IF, and CR) as the factor. The Newman-Keuls test was used as the *post hoc* test, with p < 0.05 as the significance level. Pearson’s correlation analysis was performed to evaluate associations between maternal behavioural score and SOD activity, with statistical significance set at p < 0.05. All results are expressed as mean ± standard error of the mean (SEM). Statistical analysis was performed using Statistica^®^ 10.0 software, and figures were generated in GraphPad Prism^®^ 9.0.

## 3 RESULTS

### 3.1 Nutritional assessment

A significant effect of diet (F_(2,31)_ = 68.71, p < 0.0001), time (F_(4,124)_ = 43.07, p < 0.0001), and diet × time interaction (F_(8,124)_ = 32.47, p < 0.0001) was observed for maternal body weight. Regarding the diet factor, caloric restriction dams showed the lowest body weight, followed by intermittent fasting, and then the control group. For the time factor, no significant difference was observed between PND0 and PND7, body weight increased at PND14 and PND21 and then declined by PND28. For the diet × time interaction, post hoc analyses indicated no differences among groups at PND0; however, from PND7 to PND28, intermittent fasting and caloric restriction dams exhibited lower body weight than control dams (Fig. 2A).

**Figure 2.**
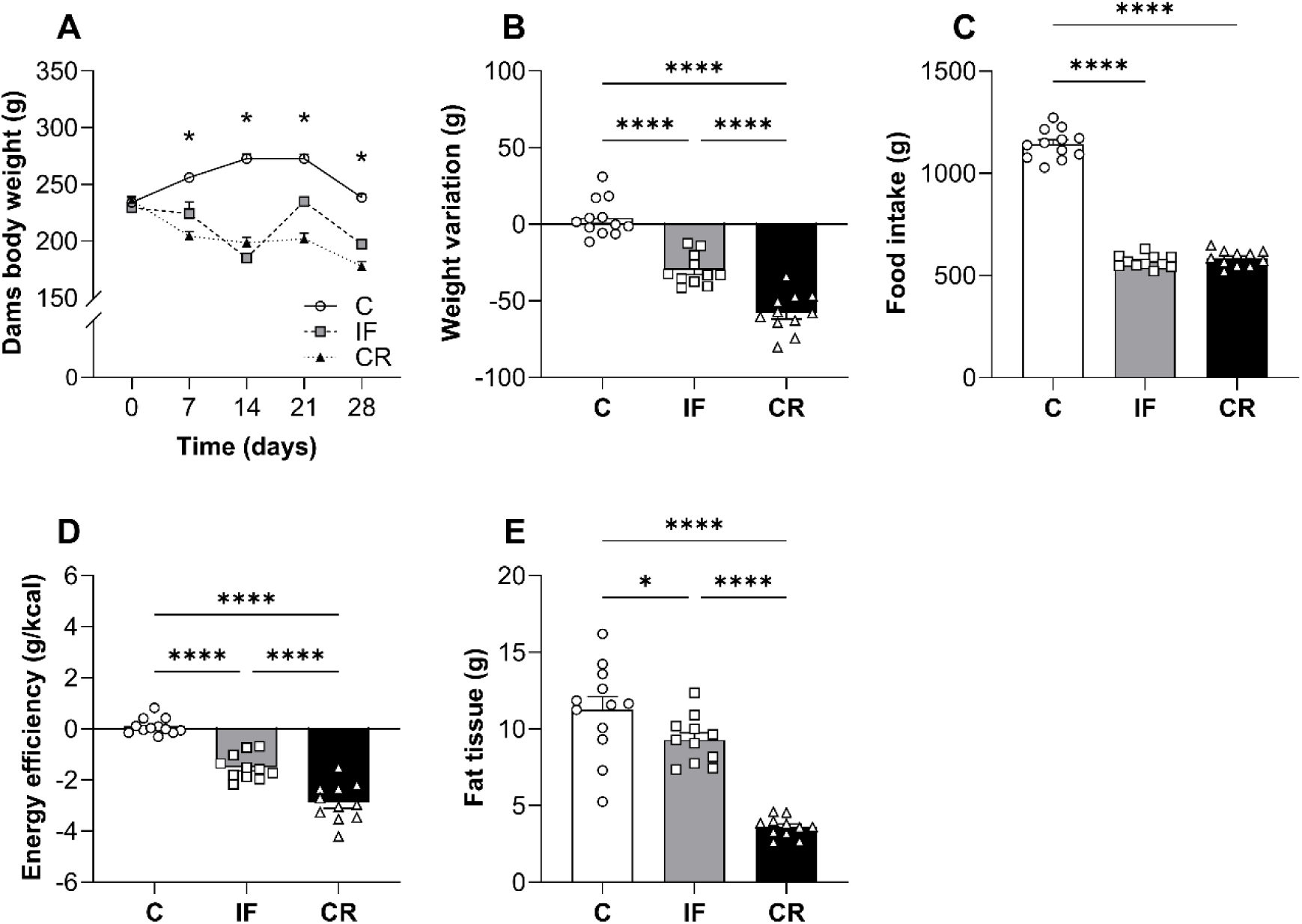
Intermittent fasting and caloric restriction in lactating dams leads to reduction in energy efficiency, body and fat tissue weights. Nutritional assessments conducted from postnatal day 0 (PND0) to PND28 in dams and litters. Body weight of dams (A); weight variation (B), total food intake (C), energy efficiency (D), and abdominal fat tissue (E) in dams. Legend: Control (C) – dams received chow *ad libitum*; IF (Intermittent Fasting) – dams feeding and fasting every 24 h; CR (Caloric Restriction) – dams received 50% of the chow consumed by Control dams. Data are shown as mean ± SEM; *n* = 11-12. \**p* < 0.05; \*\*\*\**p* < 0.0001 using ANOVA and Newman-Keuls tests.

Intermittent fasting and caloric restriction in dams had an effect on weight variation (F_(2,31)_ = 78.77, p < 0.0001; Fig. 2B), total food intake (F_(2,31)_ = 472.87, p < 0.01; Fig. 2C), energy efficiency (F_(2,31)_ = 86.13, p < 0.0001; Fig. 2D), and abdominal fat tissue weight (F_(2,30)_ = 40.92, p < 0.0001; Fig. 2E), besides BMI, body length and AC/TC ratio (Supplementary Table 1). For all these parameters, caloric restriction and intermittent fasting dams showed lower values than the control group.

Intermittent fasting and caloric restriction significantly reduced organ weights compared to the control group (Supplementary Table 2). Caloric restriction dams showed lower total cholesterol, HDL, tryglicerides, AST, creatinine, and urea blood levels than control dams; and intermittent fasting dams showed higher triglyceride levels than control dams (Supplementary Table 3).

### 3.2 Behavioural assessment

In the pup retrieval test conducted on PND6, intermittent fasting dams exhibited longer latency to retrieve the first pup (F_(2,30)_ = 5.78, p < 0.01) than control and caloric restriction dams (Fig. 3A). Intermittent fasting and caloric restriction dams showed longer latency to nest building (F_(2,30)_ = 8.17, p < 0.01) than control dams (Fig. 3B), and 72% of caloric restriction dams showed a preference for retrieving a male pup first (Fig. 3C). Intermittent fasting dams cared for their pups less time throughout the test (F_(2,30)_ = 3.50, p < 0.05) compared to control group (Fig. 3D). Intermittent fasting and caloric restriction dams displayed nursing behavior less frequent (F_(2,30)_ = 5.32, p < 0.05) and for shorter period (F_(2,30)_ = 5.79, p < 0.01) compared to control dams (Fig. 3E-F). Non-maternal behavior in the test, locomotion and rearing frequency, were similar between the three groups (Fig. 3G-H).

**Figure 3.**
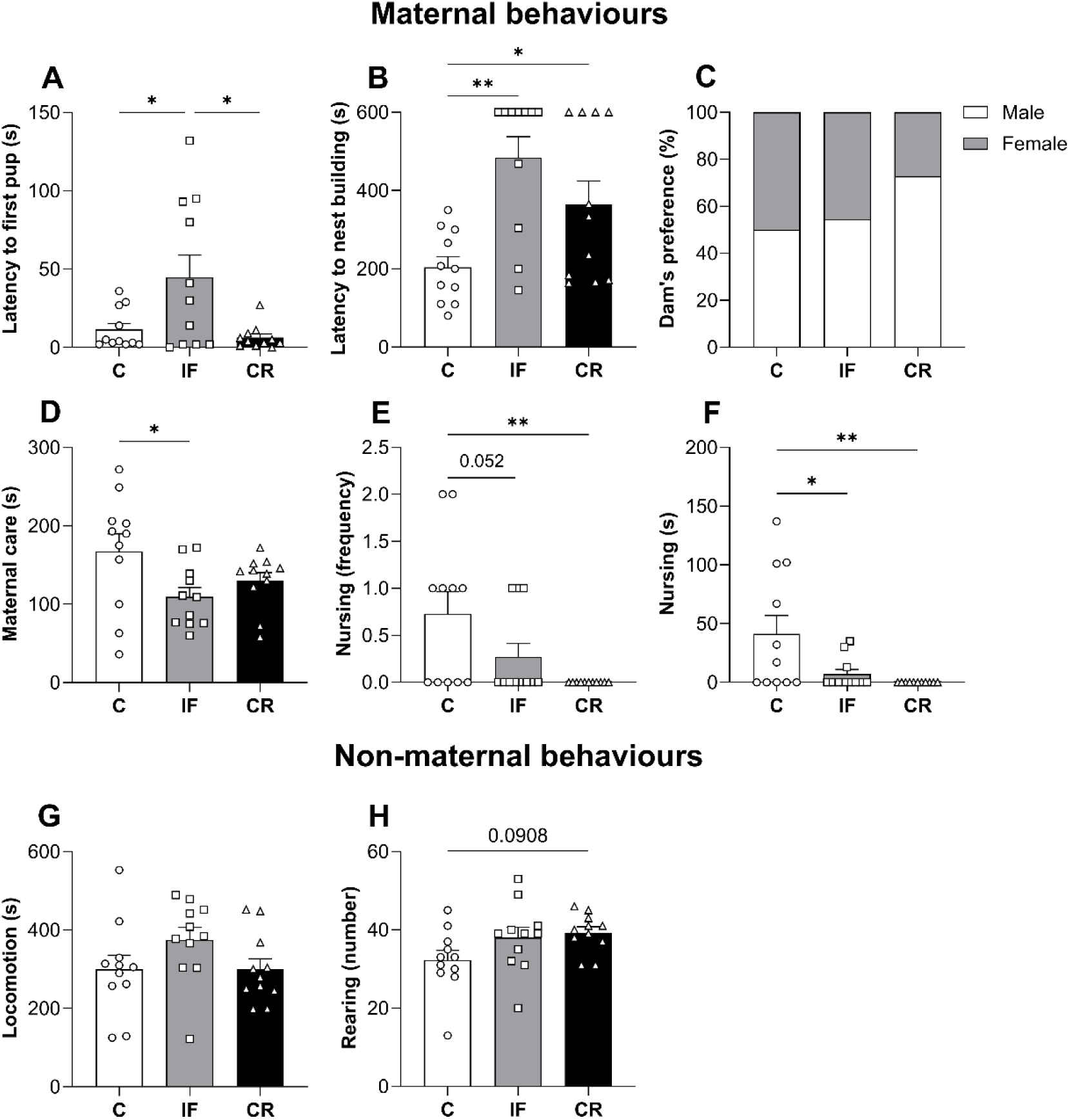
Intermittent fasting and caloric restriction in lactating dams impairs maternal behaviour on postnatal day 6. Pup retrieval test conducted on postnatal day 6 (PND6). Latency to retrieve the first pup (A); latency to nest building (B), maternal preference for retrieving a male or female pup first (C), total maternal care time (D); nursing frequency (E) and duration (F); total locomotion time (G) and rearing number (H) during the test. Legend: Control (C) – dams received chow ad libitum; IF (Intermittent Fasting) – dams feeding and fasting every 24 h; CR (Caloric Restriction) – dams received 50% of the chow consumed by C dams. Data are shown as mean ± SEM; *n* = 11. One dam (Control group) was removed from the analysis due to outlier values. \**p* < 0.05; \*\**p* < 0.01; using ANOVA and Newman-Keuls tests.

In the test conducted on PND8, intermittent fasting and caloric restriction dams exhibited longer latency to retrieve the first pup (F_(2,31)_ = 2.31, p < 0.05) than control dams (Fig. 4A). Intermittent fasting and caloric restriction dams showed longer latency to nest building (F_(2,31)_ = 8.30, p < 0.001), and 81% of caloric restrcition dams showed a preference for retrieving a male pup first (Fig. 4B). Intermittent fasting and caloric restriction dams cared for their pups less time throughout the test (F_(2,31)_ = 4.27, p < 0.05), showed lower nursing frequency (F_(2,31)_ = 8.81, p < 0.001) and duration (F_(2,31)_ = 11.23, p < 0.001), compared with control dams (Fig. 4E-F). Intermittent fasting and caloric restriction dams displayed higher locomotion (F_(2,31)_ = 5.99, p < 0.01) and rearing frequency (F_(2,31)_ = 14.05, p < 0.001), both compared to control dams (Fig. 4G-H).

**Figure 4.**
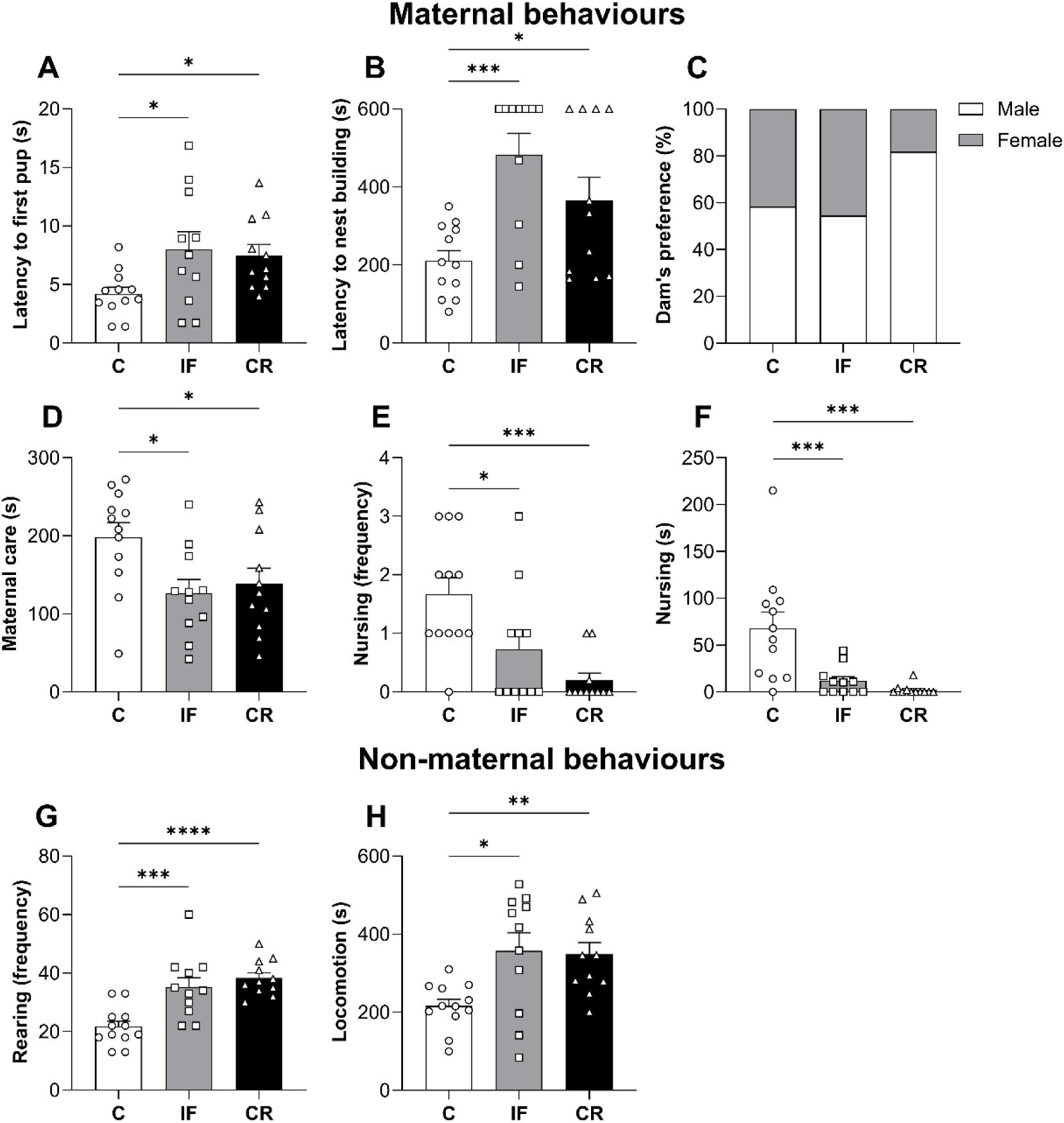
Intermittent fasting and caloric restriction in lactating dams impairs maternal behaviour and locomotion on postnatal day 8. Pup retrieval test conducted on postnatal day 8 (PND8). Latency to retrieve the first pup (A) and maternal preference for retrieving a male or female pup (B); latency to nest building (C) and total maternal care time (D); nursing frequency (E) and duration (F); total locomotion time (G) and rearing frequency (H) during the test. Legend: Control (C) – dams received chow ad libitum; IF (Intermittent Fasting) – dams feeding and fasting every 24 h; CR (Caloric Restriction) – dams received 50% of the chow consumed by C dams. Data are shown as mean ± SEM; *n* = 11-12. \**p* < 0.05; \*\**p* < 0.01; \*\*\**p* < 0.001; \*\*\*\**p* < 0.001 using ANOVA and Newman-Keuls tests.

In the EPM test, caloric restriction dams showed a lower percentage of entries in the open arms (F_(2,27)_ = 3.47, p < 0.05), compared to control dams (Fig. 5A). Caloric restriction dams showed longer time spent in the open arms (F_(2,27)_ = 7.01, p < 0.01), compared to intermittent fasting and control dams (Fig. 5B). In addition, caloric restriction dams displayed a higher frequency of visits to the distal ends of the open arms (F_(2,27)_ = 8.13, p < 0.01) than intermittent fasting and control dams (Fig. 5D).

**Figure 5.**
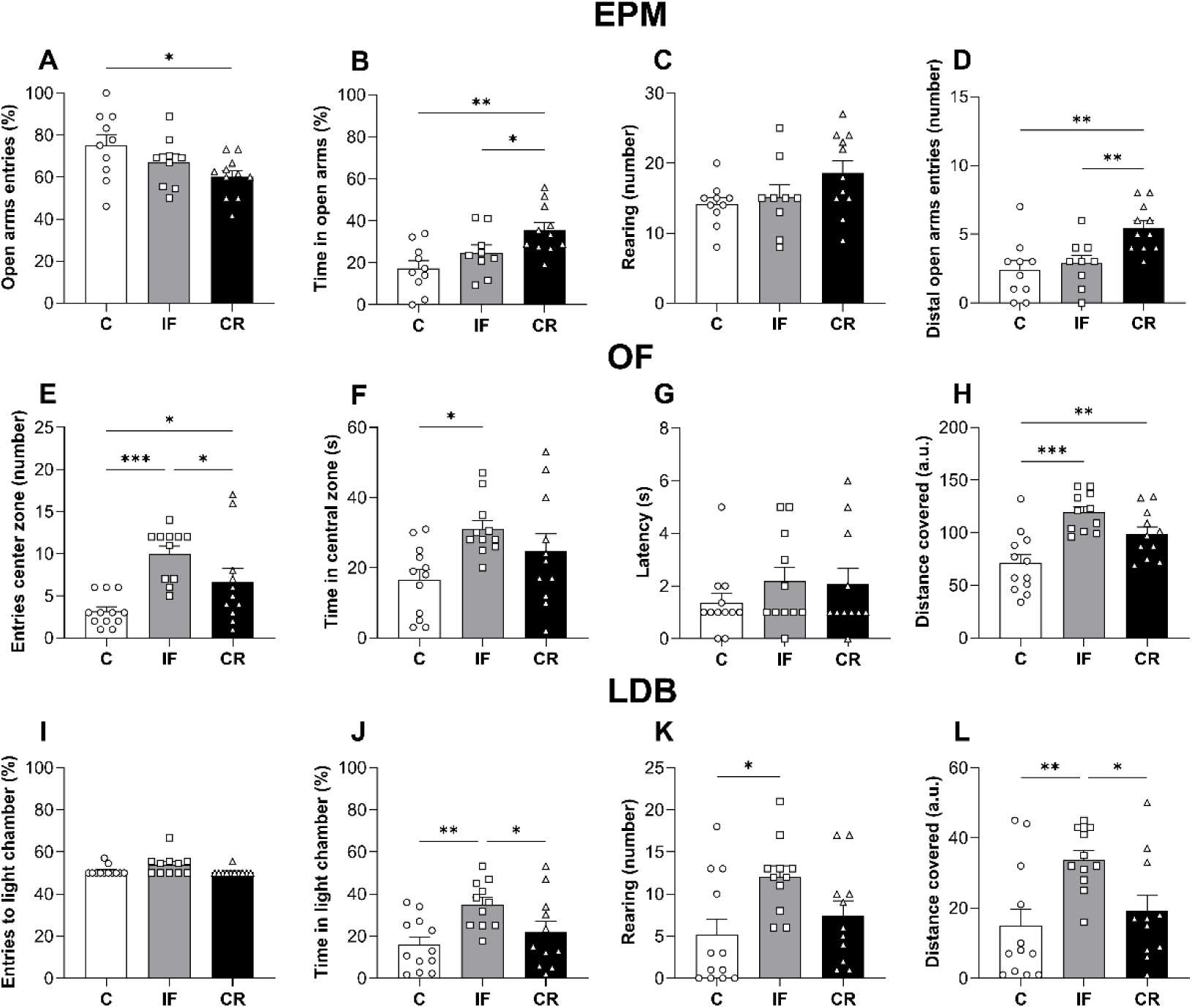
Intermittent fasting and caloric restriction promote impulsive-like behavioural profiles across distinct anxiety-related paradigms in dams. Elevated plus maze (EPM), open field (OF), and light-dark box (LDB) tests were conducted between postnatal days 22-24 (PND22-24). Percentage of entries (A) and time spent in the open arms (B), rearing frequency (C), and entries into the distal end of the open arms (D) in the EPM; entries (E), time spent (F), and latency to leave the centre zone (G), and distance travelled (H) in the OF; frequency of entries (I), time spent (J) and rearing frequency in the light chamber (K), and distance covered (L) in the LDB. Legend: Control (C) – dams received chow ad libitum; IF (Intermittent Fasting) – dams feeding and fasting every 24 h; CR (Caloric Restriction) – dams received 50% of the chow consumed by C dams. Data are shown as mean ± SEM; *n* = 11-12. EPM: Two dams (1 Control and 1 IF) were removed from the analysis due to outlier values. Two dams (1 Control and 1 IF) fell from the open arms during the test. \**p* < 0.05; \*\**p* < 0.01; \*\*\**p* < 0.01 using ANOVA and Newman-Keuls tests.

In the OF test, intermittent fasting and caloric restriction dams showed a higher frequency of entries into the centre zone (F_(2,31)_ = 10.04, p < 0.001) compared to control dams (Fig. 5I). Intermittent fasting dams exhibited a higher time spent in the centre zone (F_(2,31)_ = 4.29, p < 0.05) than control dams (Fig. 5J). Intermittent Fasting and caloric restriction dams showed a higher distance covered (F_(2,31)_ = 11.77, p < 0.001) compared to control dams (Fig. 5H).

In the LDB test, intermittent fasting dams showed a longer time spent in the light chamber (F_(2,31)_ = 5.65, p < 0.05), as well as higher rearing frequency (F_(2,32)_ = 4.30, p < 0.05) in the light chamber than control dams (Fig. 5I-J). In addition, intermittent fasting dams showed higher distance covered (F_(2,31)_ = 5.64, p < 0.01) compared to caloric restriction and control dams (Fig. 5L).

### 3.3 Redox status

Intermittent fasting and caloric restriction dams showed lower SOD activity in the amygdala (F_(2,27)_ = 12.58, p < 0.001) than control dams. In addition, intermittent fasting dams showed higher NO levels in the amygdala (F_(2,27)_ = 8.29, p < 0.01) compared to caloric restriction and control dams (Fig. 6B and D). No statistically significant differences were found for FRAP, TBARS, GST, or CAT (Fig. 6A, C and E).

**Figure 6.**
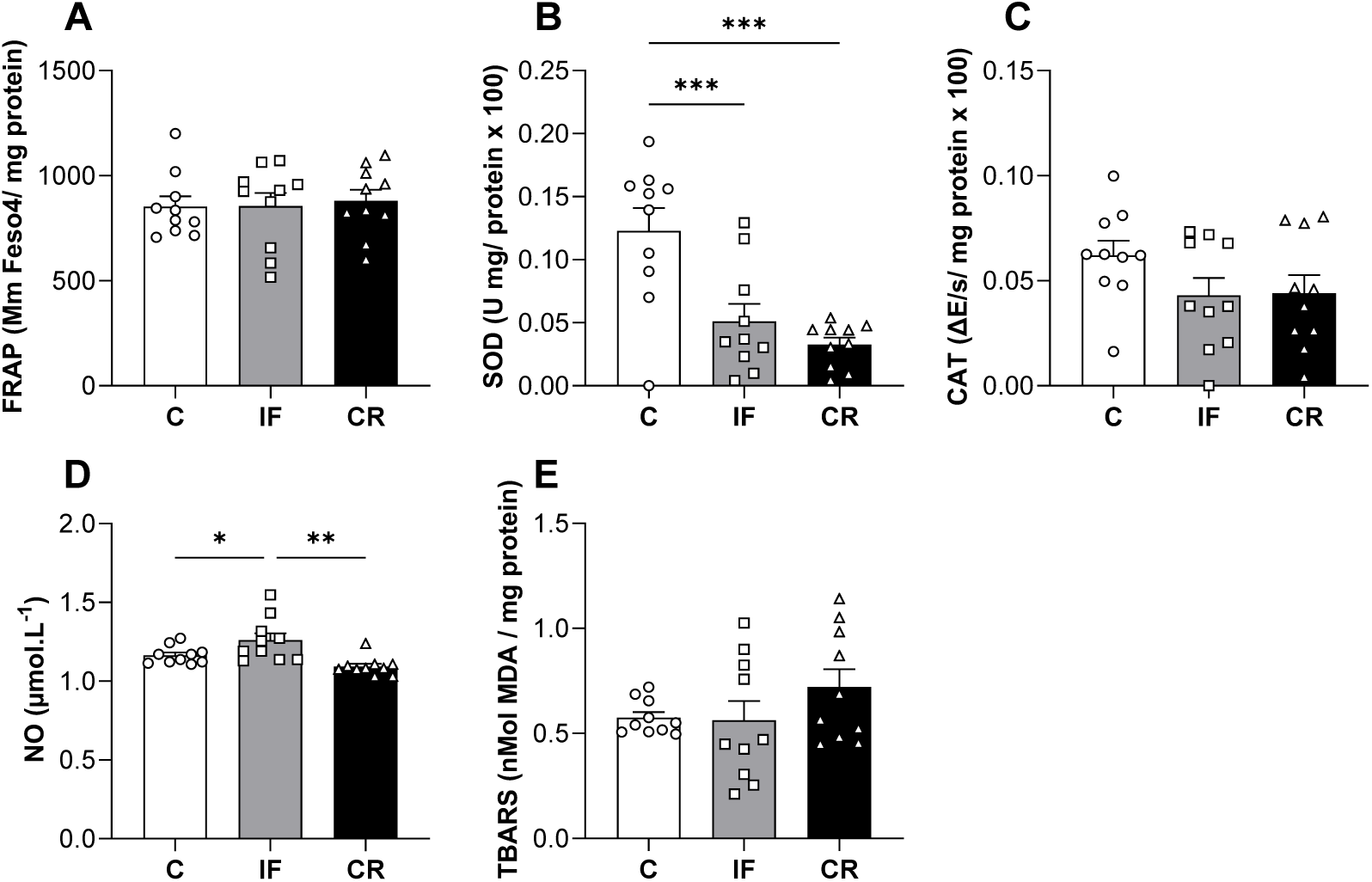
Intermittent fasting and caloric restriction reduce superoxide dismutase activity in the amygdala. Total non-enzymatic antioxidant capacity - FRAP (A); superoxide dismutase – SOD (B) and Catalase – CAT activity (C); nitric oxide (D) and lipid peroxidation – TBARS levels (E) in amygdala at postnatal day 28 (PND28). Legend: Control (C) – dams received chow ad libitum; IF (Intermittent Fasting) – dams feeding and fasting every 24 h; CR (Caloric Restriction) – dams received 50% of the chow consumed by C dams. Data are shown as mean ± SEM; *n* = 10. \**p* < 0.05; \*\**p* < 0.01; \*\*\**p* < 0.001 using ANOVA and Newman-Keuls tests.

### 3.4 Maternal behavioural score

Intermittent fasting and caloric restriction dams significantly increased the maternal behavioural score (F_(2,27)_ = 8.18, p < 0.01) compared with control group (Figure 7A). Pearson’s correlation analysis revealed a significant moderate negative association between maternal behavioural score and amygdalar SOD activity (r = -0.53, p < 0.01), indicating that higher scores were associated with lower antioxidant activity within the amygdala (Figure 7B).

**Figure 7.**
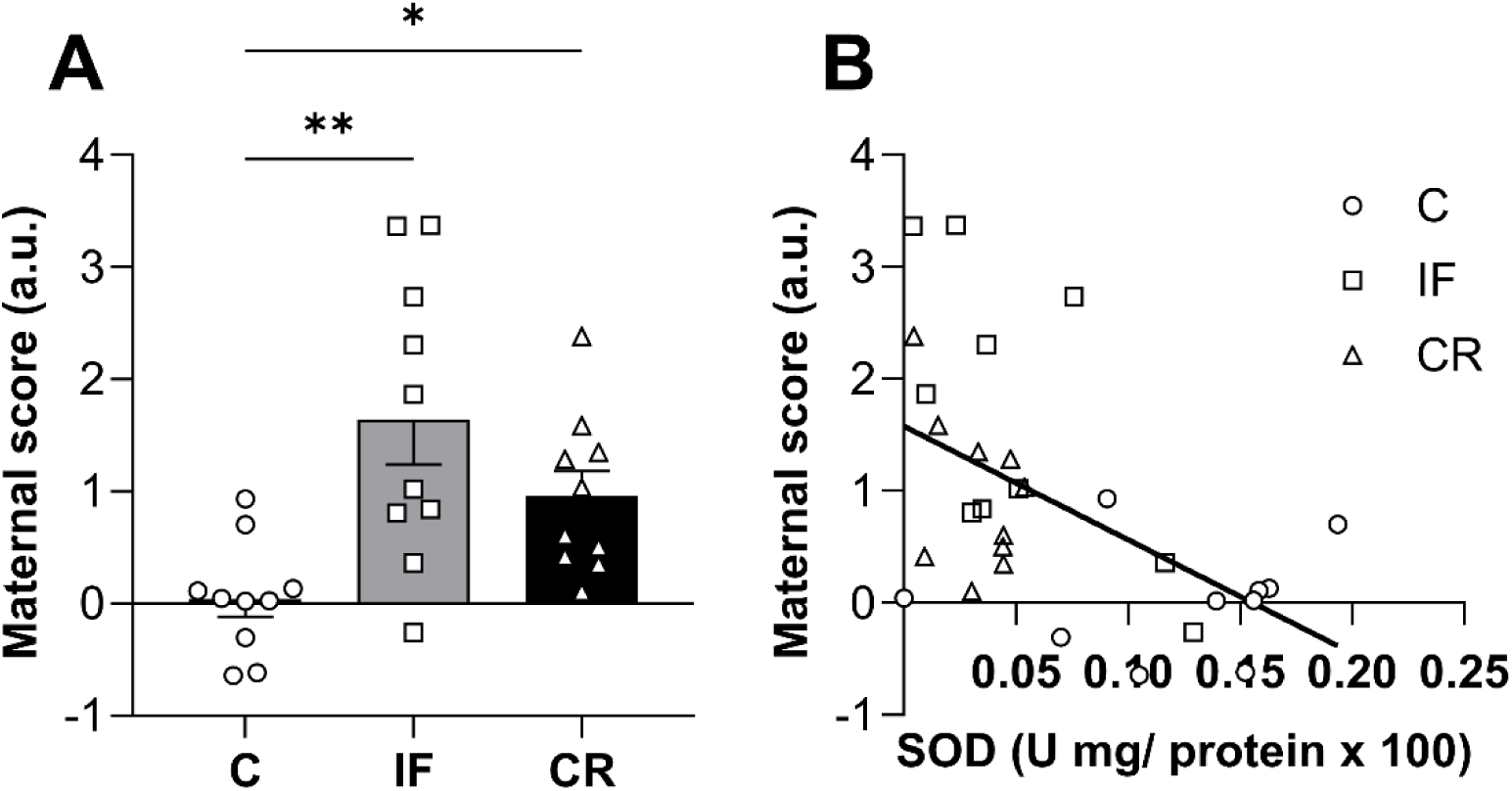
Caloric restriction increases maternal behavioural score and is negatively associated with amygdalar SOD activity. Maternal behavioural score was calculated using: first pup retrieval latency (PND6 and PND8), time spent in the open arms of the EPM, in the centre zone of the OF, and in the light chamber of the LDB (A). Pearson’s correlation analysis was performed between maternal behavioural score and SOD activity (B). Legend: Control (C) – dams received chow ad libitum; IF (Intermittent Fasting) – dams feeding and fasting every 24 h; CR (Caloric Restriction) – dams received 50% of the chow consumed by C dams. Data are shown as mean ± SEM; *n* = 10. \**p* < 0.05; ** *p* < 0.01 using ANOVA and Newman-Keuls tests.

## 4 DISCUSSION

The use of restrictive diets during critical periods, such as lactation, remains underexplored in literature with respect to maternal outcomes. Here, we employed two distinct restrictive dietary restrictions, IF and CR from PND0 to PND28, and observed that both induced underweight, impairments in maternal care quality and an increase in impulsivity. Interestingly, these behavioural alterations were negatively associated with amygdalar antioxidant activity, suggesting that higher maternal behavioural scores are linked to reduced SOD activity within this key brain region involved in emotional regulation, impulsivity, and maternal pup-directed behaviour.

Lactation can be considered a period of intense nutritional demand for the mother, encompassing her basal energy expenditure as well as the additional requirements for milk production, breastfeeding, and offspring care. Consequently, an adequate maternal diet during this phase is essential not only in terms of quantity and quality but also in feeding regularity, as irregular intake patterns are unlikely to meet the sustained metabolic demands of lactation (Da Silva et al., 2016; Muro-Valdez et al., 2023; Öner Sayar; Köseoğlu, 2025; Rocha-Gomes; Silva; Riul, 2025). In the present study, IF and CR dams exhibited marked weight loss throughout lactation, consistent with dietary models that substantially reduce daily (CR) or periodic (IF) food availability. In line with this, several other nutritional parameters were also altered in these dams, including energy efficiency – reflecting the amount of body mass gained per unit of energy intake – BMI and AC/TC ratio, as well as adipose tissue reserves. Significant reductions were also observed in circulating lipids, including total cholesterol, HDL, and triacylglycerol, directly associated with dietary restriction (Rocha-Gomes; Silva; Riul, 2025). Collectively, these findings indicate that IF and CR dams were underweight, reflecting a severe nutritional state, likely exacerbated by the metabolic demands of lactation.

Although limited, human studies indicate that undernourished mothers – particularly those experiencing food insecurity or specific nutrient deficits such as iron deficiency – are more likely to exhibit poorer parent-child relationship quality, reduced caregiving, decreased vigilance in daily tasks, and impairments in working memory (Frith et al., 2012; Prado-Silva et al., 2014). In the present study, IF and CR dams showed multiple indicators of reduced maternal care.

The pup retrieval test is a simple and widely used paradigm in preclinical rodent research to assess maternal caregiving quality and attentional engagement with the litter. It quantifies the dam’s retrieval response following pup removal from the nest, as well as a sequence of behaviours important for offspring development, including nursing and nest building (Riul et al., 1999b). Notably, both IF and CR dams showed increased latency to retrieve the first pup and reductions in key caregiving parameters, such as total maternal care time and nursing. Additionally, on PND8, IF and CR dams also exhibited increased locomotion time and rearing (vertical exploration) during the test period, non-maternal behaviours, that further support the interpretation of reduced maternal engagement and attentional focus towards the pups. Although pups become progressively less dependent on the dam as lactation advances (Riul et al., 1999b), it remained evident that, compared with Control group, IF and CR dams continued to spend less time with the litter. These findings suggest that the compromised nutritional status of these dams was associated with diminished maternal responsiveness and weakened dam-pup interactions. Importantly, disrupted early-life maternal care is also recognised risk factor for adverse mental health outcomes in offspring across the lifespan (Zhou et al., 2025).

An intriguing finding was the preferential retrieval of male pups by CR dams, with ∼70% (PND6) and ∼80% (PND8) retrieving first a male before a female. This observation suggests that under severe nutritional deprivation, such as a 50% caloric restriction, maternal behaviour may become preferentially directed towards one sex, supporting findings from other studies (Bowers et al., 2013). Previous studies have shown that male offspring exposed to maternal dietary restriction exhibit a range of adverse outcomes, including delayed pubertal onset and alterations in hypothalamic-pituitary axis function (Martins et al., 2023; Rodrigues et al., 2026). Consistent with this, our group has recently shown that CR diet during lactation induces anxiety-like behaviours in male adult offspring, whereas females exposed to the same dietary condition display milder outcomes (Rocha-Gomes et al., 2026a). Thus, we speculate that such maternal preference may contribute to the greater resilience observed in females and the heightened vulnerability in males later in life. However, further analyses and additional studies are required to support this interpretation.

Our results also indicated reduced anxiety-like behaviour, or alternatively higher impulsivity in approaching aversive spaces (Riul; Almeida, 2020), in IF and CR dams. CR group exhibited this impulsive profile in the EPM, whereas IF dams showed it in the OF and LDB tests. This divergence across paradigms may be explained by the distinct neurobiological contexts recruited by each behavioural assay. The EPM primarily involves aversion related to height and open elevation, the OF engages conflict in an open and exposed arena, and the LDB relies more strongly on visual processing and light exposure (Rocha-Gomes et al., 2021b). Thus, CR and IF diets may have induced differential neurophysiological adaptations, resulting in context-dependent responses according to the perceived type of aversion.

Additionally, our findings suggest that the reduced anxiety-like behaviour observed in IF and CR dams may be more closely related to increased impulsivity and/or hyperactivity. In a review supporting this interpretation, Pawlak and collaborators discuss that impulsiveness can be inferred from behavioural indices of risk assessment, such as approaches to the end of the open arms and increased locomotor activity (Pawlak et al., 2012). Additionally, studies in underweight individuals indicate impairments in complex cognitive processes, including decision-making and self-regulation (Lin et al., 2025), accompanied by elevated impulsivity (Lukowski et al., 2010), attention deficit (Galler et al., 2012), and is considered a risk factor for attention-deficit/hyperactivity disorder (ADHD) (Sha’ari et al., 2017). Similarly, dietary restriction models in rodents have reported increased impulsive-like responses in similar paradigms (Riul; Almeida, 2020). Together, these observations indicate that IF and CR dams not only exhibited diminished maternal engagement with their litter but also displayed heightened impulsivity, a behavioural profile that may further compromise adaptive maternal responses under challenging environmental conditions.

During lactation, dams undergo extensive neuroendocrine adaptations that prepare the brain for maternal behaviour. Multiple neural structures are recruited to support mother-pup interaction, caregiving, and nursing, among which the amygdala plays a key role. Experimental evidence indicates that pharmacological inactivation of the basolateral amygdala with muscimol reduces pup-directed contact (Numan et al., 2010), whereas optogenetic activation (Nowlan et al., 2025) enhances maternal interaction. In addition, the amygdala is intrinsically involved in anxiety-like behaviours expressed in paradigms such as the EPM, OF, and LDB (Rocha-Gomes et al., 2021b, 2026a). In this context, the redox imbalance observed in the amygdala of CR and IF dams may be linked to both the impaired performance in the pup retrieval test and the increased impulsivity observed in these mothers. The marked reduction in amino acid, vitamin, and mineral availability required for antioxidant enzyme synthesis may have contributed to decreased SOD activity observed in both IF and CR dams, alongside increased reactive nitrogen species production (nitric oxide) for IF dams (Cabrera et al., 2026; Jahoor et al., 2008; Jain; Jadhav; Varma, 2013).

Another key aspect of the present study was the calculation of the maternal behavioural score. By integrating data from the EPM, OF, and LDB tests, together with latency to retrieve the first pup (PND6 and PND8), we aimed to generate a metric reflecting maternal attention and engagement. Notably, both IF and CR dams exhibited a marked increase in this score, indicative of impaired maternal care, likely associated with the greater severity of nutritional restriction. In addition, a moderate negative correlation was observed between maternal behavioural score and amygdalar SOD activity, suggesting that reduced antioxidant activity within this brain region is associated with lower maternal attention and caregiving behaviour. To the best of our knowledge, this is the first study to link maternal behavioural impairments, increased impulsivity, and amygdalar redox imbalance in dams exposed to CR and IF during lactation.

The present study has some limitations. A more comprehensive characterisation of maternal neglect towards the litter would require continuous observation of maternal behaviour across the lactational period, as well as additional pup retrieval assessments at earlier (e.g., PND4) and later (e.g., PND12) time points in future studies. In addition, more analysis regarding the mother’s preference for one sex over the other should be conducted. The relationship between the reduced maternal engagement observed in IF and CR dams and their increased impulsivity in the EPM, OF, and LDB could be more directly examined using paradigms in which pups are placed in the open arms of the EPM, thereby allowing discrimination between pup-directed behaviour and a generalised impulsive propensity to enter aversive areas, as described by (Stolzenberg; Stevens; Rissman, 2012). Furthermore, other brain regions critically involved in maternal behaviour, such as the medial preoptic area (MPOA), as well as cortical and hippocampal regions, should be investigated in future work, including more analyses of redox status and glial cell alterations, as reported by previous studies (Dye et al., 2023; González Ibáñez et al., 2024).

## 5 CONCLUSION

In summary, our results pointed out that CR and IF diets led dams to an underweight state, impaired maternal care, and increased impulsivity, involving redox imbalance in the amygdala. From a broader perspective, this study provides a foundation for understanding the impacts of restrictive dietary regimens during the lactational period, demonstrating both nutritional and behavioural consequences for the mother. As this topic remains relatively underexplored, our findings also highlight the potential role of redox imbalance as a mechanistic contributor to the observed outcomes. Collectively, these results underscore the importance of a balanced diet, both quantitatively and qualitatively, during lactation as a cornerstone of maternal and offspring health.

## ACKNOWLEDGMENTS

This work was supported by the Coordenação de Aperfeiçoamento de Pessoal de Nível Superior – CAPES [Financial code 001]. AMS Springboard award SBF005\1102 and the MRC Career Development Award MR/T031115/1 to VM.

## DECLARATION OF AI USE

The authors used ChatGPT (OpenAI) solely to improve grammar and spelling. All scientific content, interpretations, and conclusions were developed and verified by the authors, who take full responsibility for the final version of the manuscript.

