## Supplementary table for "Caloric restriction and intermittent fasting during lactation are linked to impaired maternal care, increased impulsivity and amygdala redox imbalance in dams"

**SUPPLEMENTARY MATERIAL**

**Tabel 1**. Dam’s nutritional assessments.

|  | C | IF | CR | ANOVA |
| --- | --- | --- | --- | --- |
| Initial weight (g) | 234.43 ± 4.19 | 229.67 ± 3.20 | 235.92 ± 3,77 | NS |
| Final weight (g) | 238.61 ± 3.01^a^ | 200.00 ± 1.38^b^ | 178.13 ± 4,05^c^ | F_(2,31)_ =87.48,  p <0.0001 |
| Body lenght (cm) | 15.83 ± 0.20^a^ | 14.98 ± 0.20^b^ | 13.58 ± 0,25^c^ | F_(2,31)_ = 26.98,  p < 0.0001 |
| BMI (g/cm²) | 0.44 ± 0.004^a^ | 0.37 ± 0.005^b^ | 0.34 ± 0,013^c^ | F_(2,31)_ = 40.23,  p < 0.0001 |
| AC/TC ratio (cm) | 1.14 ± 0.013^ab^ | 1.17 ± 0.01^a^ | 1.10 ± 0,02^b^ | F_(2,31)_ = 4.43,  p <0.05 |

Nutritional assessments conducted from postnatal day 0 (PND0) to PND28 in dams and litters. Initial (PND0) and final weight of dams (PND28); body length, body mass index, abdominal/thoracic circumference ratio in dams. Legend**:** Control (C) – dams received chow *ad libitum*; IF (Intermittent Fasting) – dams feeding and fasting every 24 h; CR (Caloric Restriction) – dams received 50% of the chow consumed by Control dams. Data are shown as mean ± SEM; *n* = 11-12. Different letters within the same row indicate statistically significant differences between groups (ANOVA followed by Newman-Keuls post hoc test). Letters are assigned in descending order of the mean values, with "a" indicating the group with the highest value for the variable analysed. NS: non-significant.

**Tabel 2.** Dam’s organ weight.

|  | C | IF | CR | ANOVA |
| --- | --- | --- | --- | --- |
| Spleen (g) | 1.06 ± 0.07^a^ | 0.59 ± 0.03^b^ | 0.51 ± 0.03^b^ | F_(2,27)_ = 31.79, p < 0.0001 |
| Heart (g) | 0.84 ± 0.01^a^ | 0.65 ± 0.01^b^ | 0.58 ± 0.02^c^ | F_(2,27)_ = 18.20, p < 0.0001 |
| Liver (g) | 9.22 ± 0.51^a^ | 6.63 ± 0.14^b^ | 5.86 ± 0.40^b^ | F_(2,27)_ = 18.20, p < 0.0001 |
| Kidneys (g) | 0.97 ± 0.01^a^ | 0.88 ± 0.01^b^ | 0.81 ± 0.01^c^ | F_(2,59)_ = 118.37, p < 0.01 |
| Adrenal (g) | 0.034± 0.001^a^ | 0.025± 0.001^b^ | 0.028± 0.001^c^ | F_(2,59)_ = 27.23, p < 0.0001 |

Organs' weight was evaluated on postnatal day 28 (PND28). Legend**:** Control (C) – dams received chow *ad libitum*; IF (Intermittent Fasting) – dams feeding and fasting every 24 h; CR (Caloric Restriction) – dams received 50% of the chow consumed by Control dams. Data are shown as mean ± SEM; *n* = 11-12. Different letters within the same row indicate statistically significant differences between groups (ANOVA followed by Newman-Keuls post hoc test). Letters are assigned in descending order of the mean values, with "a" indicating the group with the highest value for the variable analysed.

**Tabel 2.** Dam’s blood assessments.

|  | C | IF | CR | ANOVA |
| --- | --- | --- | --- | --- |
| Total cholesterol(mg/dL) | 81.78 ± 2.32 ^a^ | 74.52 ± 3.79 ^a^ | 67.40 ± 2.61 ^b^ | F_(2,28)_ = 6,08,  p < 0,01 |
| HDL (mg/dL) | 40.55 ± 1.93 ^a^ | 41.31 ± 1.90 ^ab^ | 29.91 ± 3.20 ^b^ | F_(2,28)_ = 5,63,  p < 0,01 |
| Triglycerides (mg/dL) | 112.65 ± 3.77 ^b^ | 145.79 ± 3.37 ^a^ | 90.99 ± 3.57^c^ | F_(2,28)_ = 57,16,  p < 0,0001 |
| AST (mg/dL) | 71.32 ± 4.67 ^a^ | 61.63 ± 5.48^a^ | 46.18 ± 4.78 ^b^ | F_(2,28)_ = 6,54, p<0,005 |
| Creatinine (mg/dL) | 11.60 ± 1.74 ^a^ | 8.17 ± 1.10 ^ab^ | 6.89 ± 0.61 ^b^ | F_(2,28)_ = 3,69,  p < 0,05 |
| Urea (mg/dL) | 63.18 ± 4.19 ^a^ | 62.09 ± 4.32 ^a^ | 48.02 ± 1.13 ^b^ | F_(2,28)_ = 5,48,  p < 0,01 |

Blood parameters were assessed on PND28. Legend**:** Control (C) – dams received chow *ad libitum*; IF (Intermittent Fasting) – dams feeding and fasting every 24 h; CR (Caloric Restriction) – dams received 50% of the chow consumed by Control dams. Data are shown as mean ± SEM; *n* = 11-12. Different letters within the same row indicate statistically significant differences between groups (ANOVA followed by Newman-Keuls post hoc test). Letters are assigned in descending order of the mean values, with "a" indicating the group with the highest value for the variable analysed.
